# Identification of a novel SNP in the miR172 binding site of *Q* homoeolog *AP2L-D5* is associated with spike compactness and agronomic traits in wheat (*Triticum aestivum* L.)

**DOI:** 10.1101/2023.10.10.561767

**Authors:** Victoria Zeng, Cristobal Uauy, Yi Chen

## Abstract

Spike architecture is a key determinant of wheat yield, a crop which supports much of the human diet but whose yield gains are stagnating. Spike architecture mutants offer opportunities to identify genetic factors contributing to inflorescence development. Here, we investigate the locus underlying the compact spike phenotype of mutant line ANK-15 by conducting mRNA-sequencing and genetic mapping using ANK-15 and its non-compact spike near-isogenic line Novosibirskaya 67 (N67). Previous literature has placed the compact spike locus of ANK-15 to chromosome 2B. However, based on the single nucleotide polymorphisms (SNPs) identified using mRNA-seq data, we were unable to detect polymorphisms between N67 and ANK-15 in the putative chromosome 2B region. We performed differential expression analysis of developing rachis and found that *AP2L-D5*, the *D* homoeolog of the domestication *Q* gene, is upregulated in ANK-15 in comparison to N67. ANK-15 carries a SNP in the microRNA172 binding site of *AP2L-D5*, which is predicted to lead to higher expression of *AP2L-D5* due to decreased miRNA172-mediated degradation. Furthermore, we performed genetic mapping using an ANK-15 × N67 F_2_ population and found a single quantitative trait locus on chromosome 5D coinciding with the position of *AP2L-D5*. This result suggests that *AP2L-D5* is likely the underlying causal gene for the compact spike phenotype in ANK-15. We performed a field trial to investigate the effect of the *AP2L-D5* allele on agronomic traits and found that the *AP2L-D5* allele from ANK-15 is associated with a significant reduction in height, increased thousand grain weight (TGW), and increased grain width.

## Introduction

Wheat (*Triticum aestivum* L.) has played a crucial role in supporting human civilization as we rely on wheat for approximately 20% of our daily caloric and protein intake (FAO 2023). However, wheat yield increases have stagnated in recent decades, especially in major wheat-growing areas (Ray et al. 2012; Schauberger et al. 2018). Spike architecture is crucial to determine yield potential. Spike length, for example, has been shown to exhibit a high prediction ability for yield (Guo et al. 2018b). Identifying novel genetic factors regulating spike architecture can therefore provide insights into spike developmental processes and possible targets for yield improvement.

Several major loci which determine spike length and related spike morphological traits have been characterized. These include the *Q* gene on chromosome 5A and the *compactum* (*C)* locus on chromosome 2D. The dominant *C* allele is responsible for the morphology of club wheat, *T. aestivum* ssp. *compactum*, and has recently been fine-mapped to a 11 Mbp region from 231 to 242 Mb on the Chinese Spring reference genome (Kajla et al. 2023). *Q* has been molecularly characterized as a member of the *APETALA2*-like (*AP2L*) family of transcription factors (Simons et al. 2006) and is also known as *AP2L-A5* (the A subgenome homoeolog of *AP2L5*; Debernardi et al. 2020). *AP2L5* is a transcriptional repressor containing two *AP2* protein domains, and its function as a transcriptional repressor is mediated by an ethylene-responsive element binding factor-associated amphiphilic repression (EAR) motif (Liu et al. 2018). *AP2L* genes are regulated post-transcriptionally by microRNA172 (miR172), which targets *AP2L* mRNAs for cleavage (Debernardi et al. 2017). Mutations in the miR172 binding site have been shown to lead to increased *AP2L* gene expression by reducing microRNA-mediated degradation of transcripts (Greenwood et al. 2017; Xu et al. 2018). A point mutation in the miR172 target site of *AP2L-A5* led to the transition of the ancestral *q* allele, which conferred a ‘speltoid’ spike shape with an elongated rachis, to the modern *Q* allele ubiquitous in common wheat, which confers a subcompact square-shaped spike (Debernardi et al. 2017; Simons et al. 2006). In addition to controlling spike morphology, *AP2L5* is known for conferring the free-threshing character and pleiotropically influences plant height, flowering time, and grain morphology (Debernardi et al. 2022; Simons et al. 2006; Xie et al. 2018). Multiple miR172 binding site alleles of *AP2L5* have been generated with a range of phenotypic consequences. In addition to these major loci, other quantitative trait loci (QTLs) influencing spike length have been identified on several chromosomes (Liu et al. 2022; Wolde et al. 2019; Yu et al. 2022). However, few QTLs contributing to spike morphology have ultimately been cloned, suggesting that further genetic dissection is warranted.

Spike morphological mutants are useful resources for studying the mechanisms underlying inflorescence development as they represent deviations from the normal process of spike formation. To identify novel genes controlling spike length, we used near-isogenic lines (NIL) that have contrasting spike morphology. ANK-15 is near-isogenic to Novosibirskaya 67 (N67) and carries a locus that confers compact spike morphology (Koval et al. 1988). Amagai et al. (2016) showed that the compact spike locus of ANK-15 is not allelic to the *C* locus on chromosome 2D or the *Q* gene on chromosome 5A. Instead, Amagai et al. (2016) proposed the underlying gene to be allelic to a compact spike locus designated as *C_g_* from the Japanese landrace Nakate Gumbai on the long arm of chromosome 2B.

In this study, we conducted mRNA-sequencing on the developing rachis of the ANK-15 NILs in combination with genetic mapping to identify the underlying gene. To our surprise, our analysis mapped the compact spike locus onto chromosome 5D instead of the previously proposed chromosome 2B. Differential gene expression and sequence analysis support that a polymorphism on the miRNA targeted site of *AP2L-D5*, the D genome homoeolog of *Q*, is the causal mutation. Furthermore, we carried out field evaluation of the NILs to characterize the phenotypic effects of the *AP2L-D5* allele on spike morphology, plant architecture, phenology, and grain morphology.

## Results

### The *AP2L-D5* allele of ANK-15 is overexpressed and has a mutated microRNA172 (miR172) binding site

ANK-15 is near-isogenic to Novosibirskaya 67 (N67), and has a compact spike locus introduced from a mutant line of wheat cultivar Skala (Figure 1a; Amagai et al. 2016; Koval et al. 1988). Recent work suggested that the compact spike phenotype mapped to chromosome 2B (Amagai et al. 2016; Koval et al. 1988). To identify the gene underlying the ANK-15 compact spike phenotype, we conducted mRNA sequencing of the developing rachises of N67 and ANK-15 at green anther stage. We chose this stage as this is where most of the spike elongation occurs in wheat (Guo et al. 2018a). We therefore hypothesized that the causal gene for the compact spike phenotype would be differentially expressed between NILs at this stage.

**Figure 1.**
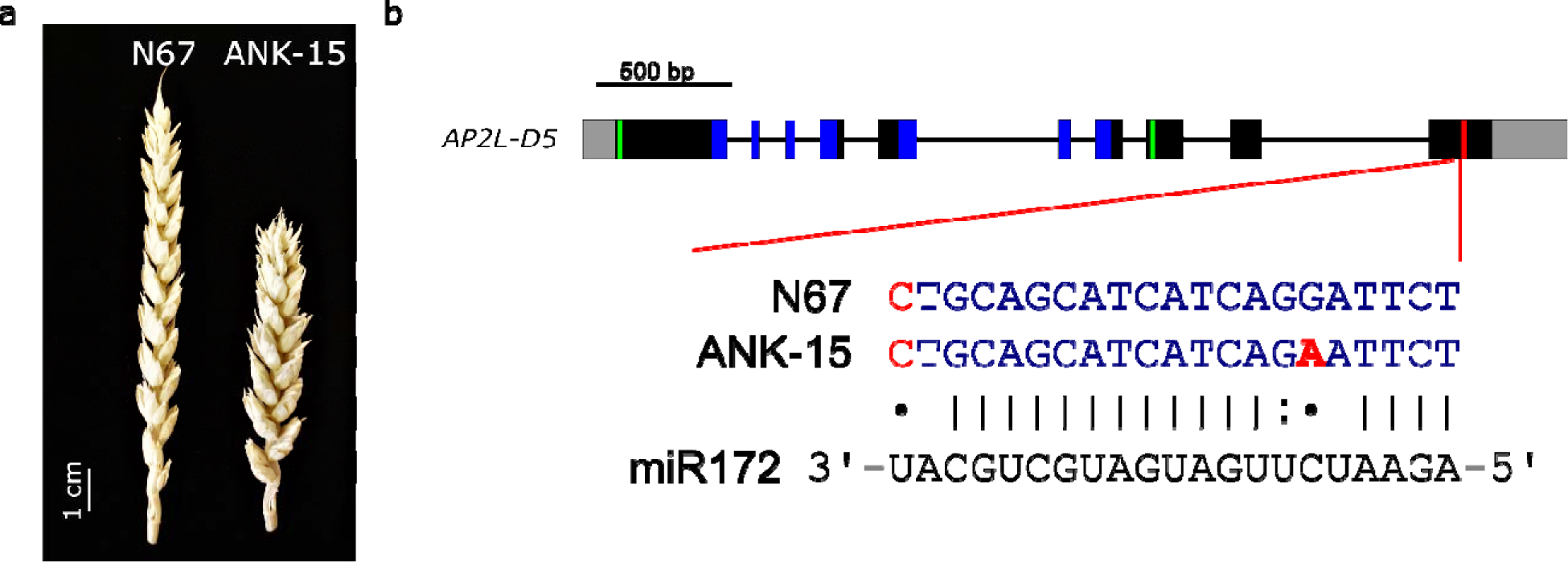
*AP2L-D5* of ANK-15 contains a single nucleotide polymorphism in the miR172 binding site. **a** Representative image of N67 and ANK-15 spikes grown in the glasshouse. **b** Intron-exon structure of *AP2L-D5* of N67 and ANK-15. Both 5’ and 3’ untranslated regions are colored in grey, exons are depicted as black boxes, introns as black lines, AP2 protein domains are highlighted in blue, EAR binding motifs are highlighted in green and the miR172 binding site is highlighted in red. Mismatches to miR172 are indicated in red font. The putative causal SNP leading to the compact spike phenotype from ANK-15 is shown in bold red font and is proposed as the *AP2L-D5b* allele.

To check the isogenic status between N67 and ANK-15, we identified polymorphisms by aligning the RNA-seq dataset against the Chinese Spring genome (RefSeq v1.0; IWGSC et al. 2018). On average, we obtained 119 million reads per sample with 91.9% of the reads aligned to RefSeqv1.0 (Supplementary table 1). As expected, we detected limited polymorphisms between ANK-15 and N67 supporting that the two lines are near-isogenic (Supplementary figure 1). There were limited polymorphisms (68 out of 1978 detected polymorphisms; 3.4%) identified on chromosome 2B. We inspected the putative interval on chromosome 2B (487 to 685 Mbp) but did not identify any polymorphisms in that interval with the closest detectable SNP about 100 Mbp away. We also identified a large polymorphic region extending across the short arm of chromosome 5A between ANK-15 and N67 (1063 out of 1978 detected polymorphisms; 53.7%).

To identify candidate genes for the compact spike phenotype of ANK-15, we conducted differential expression analysis using the RNA-seq dataset and identified 157 differentially expressed genes (DEGs) between the NILs (q < 0.05), 93 of which were high-confidence genes (Supplementary table 2). We performed GO enrichment analysis for the high-confidence gene set; however, we found no significant enrichment for any gene ontology (Supplementary table 3). To identify genes associated with spike development, we used KnetMiner, which is a gene discovery tool that infers gene function based on publicly available datasets (Hassani-Pak et al. 2021). We found that 66 out of 93 genes have functions involved in inflorescence development or are predicted to have interactions with genes of such function. Surprisingly, this includes *AP2L-D5* (*TraesCS5D02G486600*), the D subgenome homoeolog of *Q*. This gene was found to be the underlying gene of *RHT23* (*REDUCED HEIGHT 23*) and influences spike compactness and plant height (Zhao et al. 2018). This gene was upregulated in ANK-15 by 2.3-fold in the developing rachis in comparison to N67. We further performed quantitative real-time PCR (qRT-PCR) to confirm the difference in *AP2L-D5* expression between N67 and ANK-15 using samples collected from the field. We found that *AP2L-D5* was significantly upregulated (*P* < 0.05) in ANK-15 compared to N67 within the rachis (Supplementary figure 2). There was however no significant difference in relative expression of the B homoeolog (*P* = 0.37; *TraesCS5B02G486900*) and a borderline non-significant effect of the A homoeolog (*P* = 0.06; *TraesCS5A02G473800*).

Furthermore, the *AP2L-D5* allele of ANK-15 is different from N67 by a G to A transition on exon 10. We designated the wild-type *AP2L-D5* allele of N67 as *AP2L-D5a* (same as *AP2L-D5* in Chinese Spring) and the ANK-15 allele carrying the point mutation as *AP2L-D5b* based on the wheat gene nomenclature guidelines (Boden et al. 2023).

We aligned the sequences of *AP2L-D5* in N67 and ANK-15 and found that the G/A polymorphism of ANK-15 was within the miR172 binding site (Figure 1b). As mismatches to microRNA binding site are known to limit the efficiency of target mRNA cleavage, we assessed the effect of the SNP by comparing the predicted energy of interactions between miR172 and the mRNA sequences of *AP2L-D5a* and *AP2L-D5b* in RNAhybrid (Rehmsmeier et al. 2004). The minimum free energy (MFE) of the interaction between miR172 and *AP2L-D5a* was predicted to be −38.8 kcal/mol, while for *AP2L-D5b* this was −32.6 kcal/mol. The increase in the energy of interactions between miR172 and its mutated binding site on *AP2L-D5b* indicates a decrease in the binding affinity. We hypothesize that the G/A SNP in *AP2L-D5b* reduces miR172-mediated degradation of the transcript, leading to higher expression of the gene in the rachis. A different point mutation in the *AP2L-D5* miR172 binding site present in the mutant line NAUH164 (we refer to this allele as *AP2L-D5c*; Supplementary figure 3) has previously been shown to increase spike compactness in addition to pleiotropically reduce plant height (Zhao et al. 2018). Given the known connection between spike compactness and polymorphisms in the miR172 binding site of *Q*, we hypothesize that *AP2L-D5* is the underlying gene for the compact spike phenotype of ANK-15.

### The compact spike locus of ANK-15 maps to chromosome 5D not chromosome 2B

To test the hypothesis that *AP2L-D5* is the candidate gene, we performed QTL analysis using a N67 × ANK-15 F_2_ population grown in the glasshouse. As they are near-isogenic, we only developed KASP markers for three targeted regions based on the SNP data (Supplementary figure 1). First, we developed KASP marker 5D:521713197 to distinguish *AP2L-D5a* and *AP2L-D5b* in addition to other markers spanning chromosome 5D (Supplementary table 4). Second, ANK-15 was proposed to be allelic to a compact spike QTL interval (*C_g_*) from a Japanese landrace, Nakate Gumbai, between 487 and 685 Mbp on the long arm of chromosome 2B (based on BLAST location of the SSR markers; Amagai et al. 2016). However, there were limited polymorphisms on chromosome 2B (Supplementary figure 1) so we were only able to develop three KASP markers at around 770 Mbp. Lastly, we found that ANK-15 and N67 were polymorphic on chromosome 5A (Supplementary figure 1). We therefore developed markers from 176 to 539 Mbp on chromosome 5A using the identified polymorphisms.

The compact spike morphology from ANK-15 has a high density of spikelets in the apical region of the spike giving it a “squarehead” morphology (Figure 1a; Supplementary figure 4). Based on this, plants in the N67 × ANK-15 F_2_ population (*n* = 207 plants) could be visually distinguished into two distinct spike morphotypes of 140 compact spike to 67 non-compact spike lines (Supplementary figure 4). Amagai et al. (2016) considered the compact spike trait of ANK-15 to be dominant; however, a Chi-square goodness-of-fit test indicated that our observed phenotypes differed (*p* = 0.0144) from the expected 3:1 (155:52) ratio of compact to non-compact lines. The distributions of spike length, spikelet density, and height were normal in the F_2_ populations, indicating that these traits might be controlled by multiple genes in the F_2_ population.

We phenotyped the plants for spike length, spikelet density and spikelet number to perform QTL analysis. Consistent with our hypothesis, we found significant associations between markers on chromosome 5D with spike morphology and height (Figure 2b; Supplementary figure 5). The marker that distinguishes between *AP2L-D5a* and *AP2L-D5b* (5D:521713197) had the highest LOD value for spike morphological traits. F_2_ lines homozygous with the ANK-15 allele (*AP2L-D5b*; 7.9 cm) at this marker were associated with shorter spikes than F_2_ lines homozygous for the N67 allele (*AP2L-D5a*; 10.2 cm). This effect was additive with heterozygous lines showing an intermediate phenotype (9.0 cm). Marker 5D:521713197 explained 44.86% of variance in spike length. This marker is also significantly associated with spikelet number and spikelet density (Supplementary figure 5). The ANK-15 allele is associated with slightly lower number of spikelets but is still associated with higher spikelet density due to the significant reduction in spike length. The allele at *AP2L-D5* also co-segregated with the spike morphotypes. Among the 205 F_2_ plants that were successfully genotyped using marker 5D:521713197, all 56 *AP2L-D5a* homozygous lines had non-compact spikes, while all 72 *AP2L-D5b* homozygous lines had compact spikes. Of the heterozygous lines, 87 had compact spikes, while 11 were classified as non-compact. Co-segregation of the 5D:521713197 SNP with spike morphotype supports that *AP2L-D5b* is the gene underlying the ANK-15 compact spike locus. We also found that there were no significant associations with spike morphology traits for markers on chromosome 2B and 5A (Figure 2). These were the regions where the compact spike locus was proposed to be and the highly polymorphic region between N67 and ANK-15, respectively. Our genetic data thus further supports that the compact spike locus for ANK-15 is on chromosome 5D and not chromosome 2B as previously proposed, and that *AP2L-D5* is the likely causal gene.

**Figure 2.**
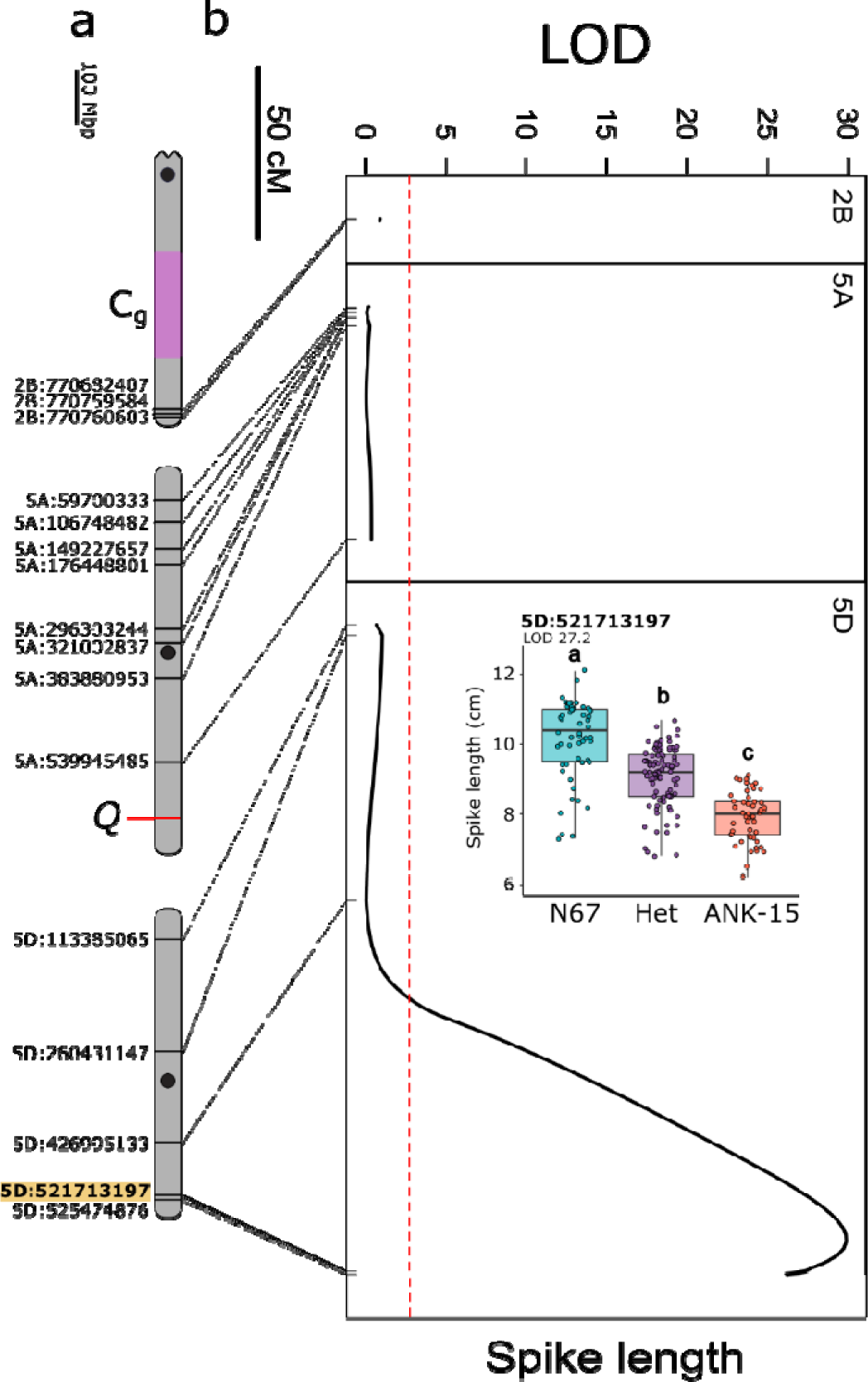
The compact spike length locus of ANK-15 maps to chromosome 5D in the N67 × ANK-15 F_2_ population. **a** Physical maps of chromosomes 2B, 5A and 5D and the KASP markers designed for genetic mapping. The markers are named after the physical position of the polymorphisms between the parents anchored onto RefSeq v1.0 (IWGSC et al. 2018). Black circles denote approximate centromere position. Putative position of the compact spike locus *C_g_*according to Amagai et al. 2016 is colored in purple. The KASP marker that separates *AP2L-D5a* and *AP2L-D5b* is highlighted in yellow (5D:521713197). **b** QTL analysis for spike length. Genetic positions of the markers are denoted with ticks and connected to their physical locations in a). The dashed red line denotes the significance threshold estimated based on permutations (*n* = 1000). **Inset** Spike length distribution of the F_2_ individuals (*n* = 207 plants) carrying parental or heterozygous alleles for KASP marker that separates *AP2L-D5a* and *AP2L-D5b*. Genotype groups significantly different from each other based on post-hoc Tukey test (*P*lJ<lJ0.01) are denoted with different letters.

Zhao et al. (2018) identified *AP2L-D5* as the candidate gene for the *REDUCED HEIGHT 23* (*RHT-23*) locus in NAUH164, an EMS mutant from Sumai 3. NAUH164 also carries a G to A point mutation within the miR172 binding site of *AP2L-D5*, although this is 10 bp upstream from the mutation that we have identified (Supplementary figure 3). The NAUH164 allele, which we proposed be called *AP2L-D5c*, led to significant reductions in plant height. We therefore hypothesize that *AP2L-D5b* can also influence plant height. In the F_2_ mapping population, we found that F_2_ lines with *AP2L-D5b* allele are associated with a significant reduction in height of 4.0 cm (Supplementary figure 5). This suggests that *AP2L-D5b* has pleiotropic effects on height in addition to spike morphology.

### *AP2L-D5b* has pleiotropic effects on agronomic traits under field conditions

The near-isogenic nature of N67 and ANK-15 was confirmed by the low number of SNPs across most chromosomes (Supplementary figure 1). Therefore, we carried out a field trial to investigate the pleiotropic effects of *AP2L-D5b* on agronomic traits, spike and grain morphology.

#### Agronomic traits

Similar to what we observed in the F_2_ population, *AP2L-D5b* is associated with a decrease in plant height under field conditions (Table 1). This decrease in plant height is in part due to a significant decrease in peduncle length. This indicated that the difference in height is not solely due to the reduction in spike length and that stem elongation is also reduced by *AP2L-D5b*. Despite ANK-15 being 5.2 cm shorter in the field, we did not observe a difference in lodging score between the NILs. We also noticed a significant effect of *AP2L-D5* alleles on heading time where *AP2L-D5b* is associated with a 2 day delay in heading.

**Table 1.** Evaluation of the pleiotropic effect of *AP2L-D5* alleles on agronomic, spike and grain morphology traits using near-isogenic lines N67 and ANK-15 under field conditions in the UK. All the traits except for height, lodging score, and heading time were measured based on five representative primary spikes/plant per plot (*n* = 6 plots). Value following the ± is the standard error of the mean. Significant effects are highlighted in red (*P* < 0.05). Thousand grain weight (TGW) was estimated using ∼ 200 grains.

| Phenotype | Lsmean |  | AP2L-D5b<br>Effect | P-value |
| --- | --- | --- | --- | --- |
|  | N67 | ANK-15 |  |  |
|  | AP2L-D5a | AP2L-D5b |  |  |
| **Agronomic traits** |  |  |  |  |
| Heading time (days) | 83.7 ± 0.2 | 85.7 ± 0.2 | 2 | 0.0001 |
| Height (cm) | 89.1 ± 0.6 | 83.9 ± 0.4 | - 5.2 | 0.002 |
| Peduncle length (cm) | 43 ± 0.6 | 39 ± 0.9 | - 4 | 0.02 |
| Stem length (cm) | 77.5 ± 0.6 | 74.6 ± 0.4 | - 2.9 | 0.02 |
| Lodging | 5.58 ± 0.7 | 6.25 ± 0.6 | 0.67 | 0.33 |
| **Spike morphology** |  |  |  |  |
| Spikelet number per spike | 20.6 ± 0.2 | 21.7 ± 0.2 | 1.1 | 0.01 |
| Spike length (cm) | 11.6 ± 0.1 | 9.3 ± 0.1 | - 2.3 | <0.0001 |
| Spikelet density | 1.78 ± 0.01 | 2.34 ± 0.03 | 0.56 | <0.0001 |
| **Grain morphology** |  |  |  |  |
| Number of grains per spike | 59.2 ± 1.1 | 53.6 ± 1.0 | - 5.6 | 0.02 |
| Thousand grain weight | 41.2 ± 0.5 | 44.1 ± 1.0 | 2.9 | 0.02 |
| Grain area (mm<sup>2</sup>) | 17 ± 0.1 | 17.6 ± 0.3 | 0.6 | 0.05 |
| Grain length (mm) | 6.32 ± 0.01 | 6.42 ± 0.05 | 0.1 | 0.17 |
| Grain width (mm) | 3.32 ± 0.02 | 3.41 ± 0.03 | 0.09 | 0.02 |

#### Spike morphology

Similar to what was observed in the F_2_ population grown in the glasshouse, *AP2L-D5* alleles have significant effects on spike length and spikelet density. *AP2L-D5b* in ANK-15 was associated with a decrease in spike length and increase in spikelet density (Table 1). Interestingly, *AP2L-D5b* was associated with an increase in spikelet number under field conditions but was associated with a decrease in spikelet number under our glasshouse conditions (Table 1, Supplementary figure 5).

#### Grain morphology

*AP2L-D5* has a significant impact on grain width where *AP2L-D5b* is associated with wider grains than *AP2L-D5a* (Figure 3; Table 1). This effect however was not observed for grain length (*P* = 0.17). The increase in grain width translated to an increase in thousand grain weight (TGW; *P* = 0.02; Table 1) in the *AP2L-D5b* NILs. However, the increase in TGW is compensated with a decrease in the number of grains per spike as measured on the primary spike. Overall, *AP2L-D5* alleles, in addition to affecting spike morphology, have pleiotropic effects on agronomic traits and grain morphology.

**Figure 3.**
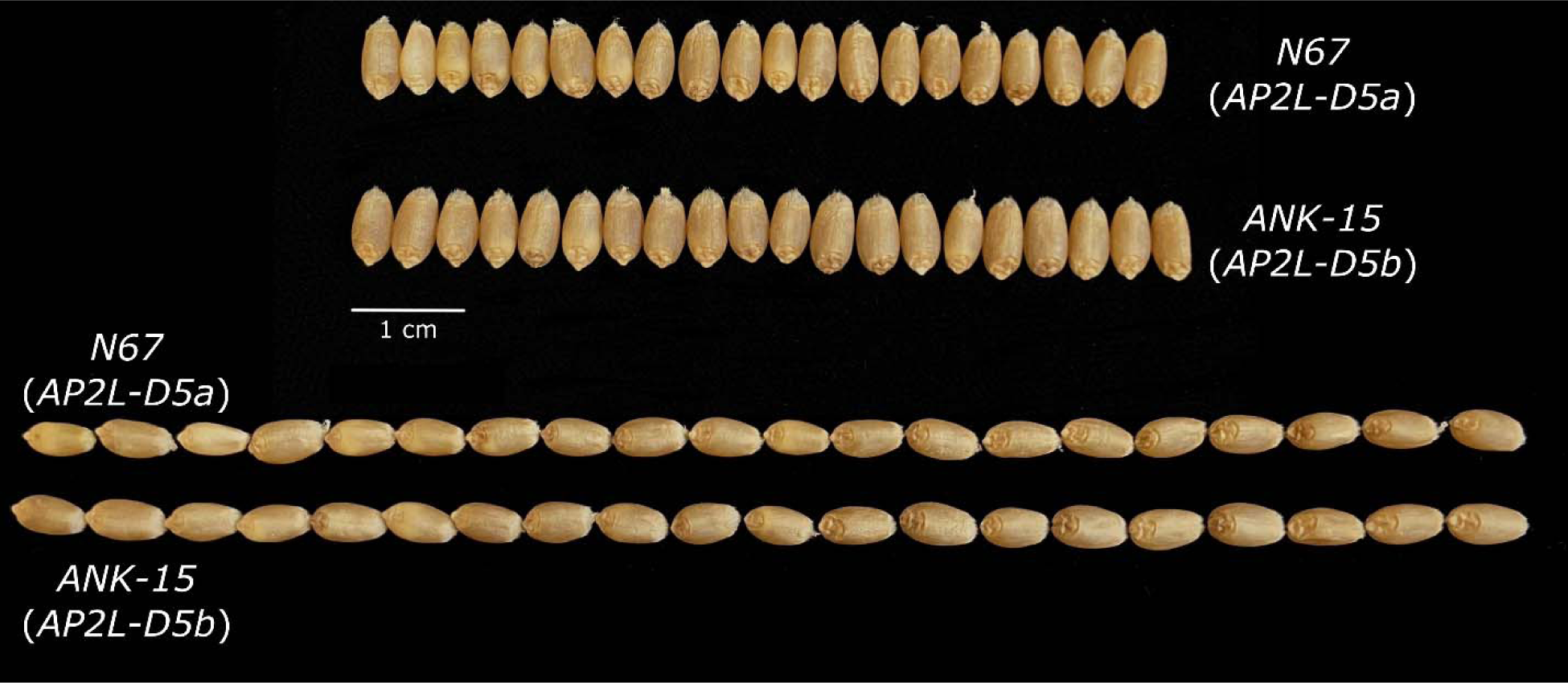
Representative grains of N67 and ANK-15 to illustrate the effect of *AP2L-D5* alleles on grain morphology.

## Discussion

Identifying the genes involved in spike development can reveal targets for yield improvement. In this study, we identified an allele of *AP2L-D5* with a novel SNP in the miR172 binding site. Similar to the mutation in *AP2L-A5* that gave rise to the *Q* allele, the SNP in *AP2L-D5b* leads to increased expression which we attribute to reduced transcript degradation due to the expected decreased binding affinity for miR172. Several natural and induced miR172-resistant mutants have been identified for *Q* (Debernardi et al. 2017; Greenwood et al. 2017; Liu et al. 2020; Xu et al. 2018). However, only one miR172 binding site mutant, NAUH164, has been previously reported for *AP2L-D5* (Zhao et al. 2018). Our results provide greater insight into the role of miR172-mediated regulation of *AP2L-D5* as well as the possible breeding utility of this novel allele.

ANK-15 carries a compact spike locus introduced from a mutant line of wheat cultivar Skala. Recent work on ANK-15 proposed that the locus is allelic to a novel compact spike locus, *C_g_*, from the Japanese landrace Nakate Gumbai (Amagai et al. 2016). However, we did not detect any polymorphisms in the putative region between N67 and ANK-15, which are near-isogeneic lines developed using phenotypic selection (Koval et al. 1988; Supplementary figure 1). In addition, genetic mapping of the F_2_ population of N67 × ANK-15 found no significant association between the markers on chromosome 2B and spike morphology (Figure 2b). Instead, we found significant association between the markers on chromosome 5D with spike morphology, showing that the compact spike locus of ANK-15 is on chromosome 5D. By using mRNA-seq, we found that *AP2L-D5* is expressed at a higher level in ANK-15 in comparison to N67 (Supplementary table 2; Supplementary figure 2). ANK-15 carries a point mutation on the miR172 binding site of *AP2L-D5* which introduces a mismatch between miR172 and its binding site (Figure 1). Study has shown that mismatch between miR172 and its binding site limit its cleave efficiency resulting in higher transcript level of its target (Debernardi et al. 2017) which we have observed in ANK-15 in both the RNA-seq and qRT-PCR (Supplementary figure 2). A different mutation that introduced a mismatch in the miR172 binding site of *AP2L-D5* (*AP2L-D5c*; Supplementary figure 3) also led to compact spike (Zhao et al. 2018) and the same mutation as ANK-15 but on the A subgenome homoeolog (*Q* gene) also led to compact spike morphology (Greenwood et al. 2017). *AP2L-D5b* has a dosage-dependent effect on spike length and height (Supplementary figure 5). The *AP2L-D5c* allele of NAUH164 is similarly additive, as heterozygous F_2_ lines were intermediate for these traits (Zhao et al. 2018). Similarly, in the F_2_ progeny of a cross between a miR172 binding site mutant of *AP2L-A5* and its wild-type cv. Sunstate, Greenwood et al. (2017) observed that heterozygotes had intermediate spike compactness and height. Based on the strong connection between *AP2L5* genes and spike morphology, we propose that *AP2L-D5* is likely the underlying gene of the compact spike morphology of ANK-15.

As the wheat spike is determinate, spikelet number per spike (SNS) is limited early in development by the number of axillary meristems formed before the terminal spikelet stage. Delaying the terminal spikelet stage may therefore lead to higher spikelet numbers, albeit often with an associated delay in flowering time. Greenwood et al. (2017) found an increase in rachis node number (which are the nodes that individual spikelets are attached to) with increased *AP2L-A5* expression. Conversely, SNS decreased in *ap2l5* null mutants generated by Debernardi et al. (2020) and in a *Q* truncation mutant identified by Zhang et al. (2022). These results indicate that *AP2L5* can delays formation of the terminal spikelet leading to higher SNS. Consistent with these results, our field-grown ANK-15 plants with increase *AP2L-D5* expression exhibited greater number of spikelets in comparison with N67 (Table 1). However, *AP2L-D5b* was associated with a decrease in SNS in F_2_ lines grown in the glasshouse. These discrepancies may be explained by differences between glasshouse and field conditions. Uniform temperatures and photoperiod in the glasshouse may have promoted greater uniformity and faster floral development across the F_2_ genotypes.

A delay in terminal spikelet formation is one component of altered phenology caused by increased *AP2L5* expression. The *ap2l5* null mutants developed by Debernardi et al. (2020) as well as transgenic miR172 overexpression lines (Debernardi et al. 2017) flowered earlier than their wild-type lines, which is reflected in our field evaluation by the delayed heading exhibited by ANK-15 (Table 1). Through differential expression analysis, we found that *TERMINAL FLOWER1* (*TFL1*; *TRAESCS5A02G128600*), a floral repressor, was upregulated in ANK-15 during spike development, suggesting it may be regulated (directly or indirectly) by *AP2L-D5* (Supplementary table 2). *TFL1*, a phosphatidylethanolamine-binding protein, regulates inflorescence meristem identity and is a known negative regulator of flowering time in *Arabidopsis* (Simon et al. 1996as). Yant et al. (2010) demonstrated that *TFL1* was downregulated in *ap2* mutants. The transcription factor *AP1*, a target of *AP2*, directly represses *TFL1* activity (Kaufmann et al. 2010), which may underlie the upregulation of *TFL1* in *AP2* family overexpression mutants such as ANK-15. However, the association between *TFL1* and *AP2L5* expression in wheat has not been characterized to date. Our results suggests that it would be of interest to further explore if *AP2L5* regulates meristem transition through *TFL1*.

Although an increase in spikelet number due to delayed terminal spikelet formation may be expected to increase grain number per spike, our grain morphometric analysis revealed that ANK-15 spikes had a significantly lower average grain number per spike compared to N67 (Table 1). These results suggest that *AP2L-D5* might influence the distribution of yield components, as the reduction in grain number corresponded to an increase in TGW and grain width (Table 1; Figure 3). Our ANOVA results also suggest that *AP2L-D5b* may positively influence grain area. Xie et al. (2018) demonstrated that the transformation from *q* to *Q* caused by a miR172 binding site SNP, led to increased grain width and TGW. Larger grain size is beneficial not only from a yield perspective but is also associated with increased early vigor (Zhao et al. 2019). In addition, milling yield is positively correlated with grain size (Marshall et al. 1986). Hence, the enhancement of grain width represents a potential way in which this novel allele may contribute to variety improvement. Interestingly, the miR172-resistant *AP2L-D5* allele of NAUH164 conferred a reduction in both TGW and grain number per spike (Chen et al. 2015; Zhao et al. 2018), suggesting that different polymorphisms in the miR172 binding site may differ in their pleiotropic effects on yield and quality-related traits. The difference between the studies can also be attributed to the difference in genetic background and environment in which these studies were conducted.

Beyond inflorescence development and grain morphology, *AP2L-D5b* also has pleiotropic effects on plant height (Table 1). The mechanism by which *AP2L5* genes affect height has not been fully characterized. Patil et al. (2019) proposed that miR172-mediated *HvAP2* degradation in barley ultimately facilitates gibberellic acid (GA)-promoted stem elongation, finding that a miR172-resistant mutant showed inhibited growth responses to GA. Several semi-dwarfing loci in use since the Green Revolution, such as *Rht-B1b* and *Rht-D1b*, encode truncated DELLA proteins that constitutively repress GA responses (Thomas 2017; Van De Velde et al. 2021). The use of these GA-insensitive semi-dwarfing alleles leads to reduction in internode elongation, but the disruption of the GA signal transduction pathway also confers undesirable pleiotropic effects on seedling emergence and growth due to coleoptile shortening (Allan 1980; Tang et al. 2009). Therefore, alternative semi-dwarfing genes are currently sought after (Borrill et al. 2022). While exogenous application of GA to the miR172-resistant NAUH164 does not lead to the restoration of wild-type height, an alpha amylase assay demonstrated that the GA signaling pathway was not disrupted in the mutant at the seed stage (Chen et al. 2015). This may indicate that GA insensitivity caused by *AP2L5* overexpression could be constrained to the stem elongation process. Although further insight into the relationship between AP2 genes and the GA pathway is needed, miR172-resistant alleles such as *AP2L-D5b* may present alternative sources of height reduction that avoid the limitations of the *Rht1* alleles caused by constitutive GA insensitivity. The effect of *AP2L5* alleles on spikelet density might be a concern for mechanical harvest but production of club wheat with similar spike structure, to the best of our knowledge, has not been linked with harvest issues.

## Materials and methods

### Plant materials and growth conditions

We developed an F_2_ mapping population derived from crossing near-isogenic lines ANK-15 and N67 (*n* = 207). Seeds were stratified at 4°C for 48 hours, then sown in John Innes Cereal Mix (65% peat, 25% loam,10% grit, 3 kg/m^3^ dolomitic limestone, 1.3 kg/ m^3^ PG mix (14-16-18; Yara UK), 3 kg/ m^3^ Osmocote Exact) in glasshouses at the John Innes Centre, Norwich, UK. Plants were grown under long-day (16-h light: 8-h dark) conditions at 21 °C and 16 °C during the day and night, respectively. Alongside the ANK-15 F_2_ population, we also grew the parental varieties N67 (*n* = 6 plants) and ANK-15 (*n* = 5 plants) for phenotypic comparison with the F_2_ progenies.

We grew N67 and ANK-15 at the Dorothea de Winton field station in Bawburgh, UK (52°37′50.7”N 1°10′39.7”E) during the 2023 growing season (sown 27 March 2023) to further characterize the phenotypic effects of the allele under field conditions. Each NIL was drilled in 1.3 m^2^ (1 m x 1.3 m) double-row plots with a row spacing of 0.17 m in randomized complete block design with six blocks. Plant growth regulator (3C Chlormequat 750; BASF) was applied to all plots 53 days after sowing at the rate of 0.6 L/ha.

### Phenotyping

To phenotype the N67 × ANK-15 F_2_ population, we measured the spike length (SL), spikelet density (SD), spikelet number per spike (SNS), and plant height at maturity for the primary spike. We also visually scored plants as having either a non-compact (wild-type) or compact (ANK-15-like) spike morphotype. SL is the distance between the first basal spikelet node to the tip of the spike, excluding awns. SD is calculated as 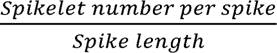. Plant height was measured as the distance between the soil and the tip of the tallest culm without awns.

In our field evaluation of the *AP2L-D5* NILs, heading date was the number of days at which 75% of spikes in a plot had emerged. We assigned a lodging score to each plot at the end of the growing period based on lodging incidence, in which a score of 1 indicated high lodging and 9 indicated no lodging. Five representative primary spikes were selected for phenotyping SL, SNS, and SD in each of the six plots. In addition, we recorded architectural traits including the lengths of the primary tiller (plant height), stem (distance from base of culm to base of spike), and peduncle (base of spike to last node on stem). To assess the effects of the compact spike trait on grain morphology, we used a MARVIN grain analyzer (GTA Sensorik GmbH, Germany) to record average grain morphometric traits, including grain number and grain weight per spike, grain width, length, area, and thousand grain weight (TGW), of the five subsamples per plot.

### Rachis transcriptome analysis of ANK-15 and N67

We collected and froze the rachises of four ANK-15 and four N67 spikes (one rachis per one plant) at green anther stage (Guo et al. 2018b) in liquid nitrogen and homogenized individual samples with a mortar and pestle. Using the Spectrum Plant Total RNA kit (Sigma), we isolated RNA from each sample according to the manufacturer’s protocol and performed On-Column DNAse I digestion (Sigma) to remove contaminating genomic DNA. To quantify the RNA concentration of each preparation, we used a NanoDrop 1000 spectrophotometer (Thermo Fisher). The mRNA-sequencing was performed by Novogene (Cambridge, UK) using the NovaSeq 6000 platform (Illumina, United States), generating paired-end 150 bp reads. We aligned these reads to the Chinese Spring genome RefSeq v1.0 assembly (IWGSC et al. 2018) using HiSAT2 (v2.1.0; Kim et al. 2019). Using SAMtools (v1.4.1; Danecek et al. 2021), the resulting alignments were converted to BAM files, sorted, and indexed, allowing visualization in the Integrative Genomics Viewer browser (IGV; Robinson et al. 2011).

To identify differentially expressed genes (DEGs) between the NILs, we quantified the abundance of transcripts with Kallisto (v0.48.0), using 30 bootstrap samples (Bray et al. 2016). Reads were pseudoaligned to a transcriptome index constructed using the IWGSC RefSeqv1.0 assembly and v1.1 annotation. Differential analysis of the quantified transcripts was then performed using the R package Sleuth (v0.30.1; Pimentel et al. 2017). We identified DEGs that function in infloresence development by using the web-based gene function discovery platform KnetMiner with the search term “inflorescence” (Hassani-Pak et al. 2021). Gene ontology (GO) terms of DEGs were extracted with Ensembl BioMart (Kinsella et al. 2011) and analyzed in the PANTHER GO enrichment analysis tool on the Gene Ontology Resource webpage (Mi et al. 2013).

### Genotyping

To identify polymorphisms between parental lines, we called variants from RNA-seq alignments using FreeBayes (v1.3.1; Garrison and Marth 2012) with a minimum alternate allele count of 5. We retained SNPs with a quality score of at least 1000 and a read depth of at least 40. SNPs on candidate chromosomes were selected for conversion to Kompetitive Allele-Specific PCR (KASP) markers using the PolyMarker (Ramirez-Gonzalez et al. 2015) primer design pipeline. The 5’ termini of allele-specific forward primers were labeled with the FAM (5′-GAAGGTGACCAAGTTCATGCT-3′) or HEX (5′-GAAGGTCGGAGTCAACGGATT-3′) tail sequence. All primers specific to SNPs from N67 were FAM-compatible (Supplementary table 4).

To genotype the N67 × ANK-15 F_2_ population, we conducted KASP assays in a 384-well format. For each marker, we prepared a primer assay mix consisting of both allele-specific forward primers (12 μM) and the common reverse primer (30 μM) (Supplementary table 4). Each reaction consisted of 2 μL DNA (10-50 ng), 1.944 μL PACE Genotyping Master Mix (Standard ROX; 3CR Bioscience), and 0.056 μL primer assay mix. PCR was carried out according to the following conditions: initial denaturation and hot-start enzyme activation at 94 °C for 15 min; 10 touchdown cycles of template denaturation (94 °C, 20 s) followed by annealing/extension with decrease of 0.8 °C per cycle (65–57 °C, 60 s); and 40 cycles of amplification consisting of a denaturation step (94 °C, 20 s) followed by annealing/extension (57°C, 60 s). Fluorescence of PCR products was detected with a PheraStar plate reader (BMG Labtech, Germany). We identified genotypes manually in KlusterCaller (version 3.4.1; LGC Biosearch Technologies, UK). Samples showing ambiguous clustering were subject to additional 5-cycle amplifications (94 °C, 20 s; 57 °C, 60 s) to improve separation of clusters.

### Sequencing of the candidate gene *AP2L-D5*

After identifying *AP2L-D5* as a possible candidate gene, we amplified its sequence in ANK-15 and N67 (Supplementary table 5). PCR was carried out according to the conditions specified in the source publication of the primer sequences (Zhao et al. 2018). PCR products were purified using the Wizard SV Gel and PCR Clean-Up System (Promega) in line with the manufacturer’s protocol. Samples were sequenced using the amplicon sequencing service provided by Plasmidsaurus which used Oxford Nanopore sequencing.

### Energy prediction of miR172-target interaction

To predict the binding affinities of the wild-type and mutant *AP2L-D5* alleles with miR172, we used the RNAhybrid online web tool (Rehmsmeier et al. 2004) which computes the minimum free energy (MFE) hybridization of a pair of RNA molecules. The mRNA sequences of *AP2L-D5* from N67 and ANK-15 and the sequence of wheat miR172 (Debernardi et al. 2017) were uploaded to the RNAhybrid server.

### Quantitative Reverse Transcription PCR of candidate genes

To verify the results from RNA-seq, we compared the expression levels of the candidate gene and its homoeologs in rachises collected at yellow anther stage between NILs grown in the field (Guo et al. 2018b). Four biological replicates, consisting of one rachis from each of the four plots, were sampled from each genotype. We extracted and quantified RNA as above and synthesized first-strand complementary DNA (cDNA) from 1 μg of total RNA per sample using the SuperScript III First-Strand Synthesis System (Invitrogen). Each reaction consisted of 5 μL LightCycler 480 SYBR Green Master mix (Roche), 0.5 μL each of the forward and reverse primers (100 μM; Supplementary table 5), 3 μL nuclease-free water, and 1 μL of diluted cDNA (1:5) for a total volume of 10 μL. *Glyceraldehyde-3-phosphate dehydrogenase* (*GAPDH*) was selected as an endogenous reference gene for expression normalization.

For qRT-PCR, we performed three technical replicates and three no template control (NTC) reactions for each biological replicate. qRT-PCR was conducted on a LightCycler 480 Instrument II (Roche) under the following conditions: initial template denaturation at 95 °C (5 min); 45 amplification cycles, each consisting of denaturation at 95 °C (10 s), annealing at 62 °C (15 s), and extension at 72 °C (30 s); generation of a dissociation curve from 60 °C to 95 °C. Upon completion of the qRT-PCR, we verified that the amplified products were target-specific through melting curve analysis. To analyze the relative expression levels of the target genes, we employed the 2^−ΔΔCT^ method (Livak and Schmittgen 2001) and compared gene expression between N67 and ANK-15 using Student’s t-tests.

### QTL mapping and statistical analysis

QTL analyses were conducted using the *R/qtl* package (v1.60; Broman et al. 2003) in R (v4.3.0; R Core Team, 2023). We calculated conditional genotype probabilities and marker recombination frequencies in the F_2_ populations, following Haldane’s mapping function. Assuming a single QTL, we conducted standard interval mapping using the *scanone* function with the expectation-maximization algorithm. We employed the normal model for data with a normal distribution and the non-parametric model for data that failed the Shapiro-Wilk test.

Statistical analyses of phenotypic data were performed in R. To determine whether parametric statistical tests were appropriate, we conducted Shapiro-Wilk tests to assess the normality of phenotypic data. We conducted one-way analysis of variance (ANOVA) tests to evaluate the effect of *AP2L-D5* alleles on the traits measured. Tukey’s Honestly Significant Difference (HSD) test was implemented to further identify pairwise differences between the genotypes.

For the field evaluation of *AP2L-D5b*, we assessed the phenotypic effects of the genotype and the blocking factor (plot) by aggregating the five subsamples taken from each plot to obtain a plot mean for each phenotype and performed two-way ANOVAs including genotype and block as factors. To evaluate the effect of genotype and block when a non-parametric test was required, we conducted permutational two-way ANOVA based on 5000 permutations using the *aovp()* function from the R package *lmPerm* (v2.1.0; Wheeler and Torchiano 2016).

## Statements & Declarations

### Author Contribution

Conceptualization: YC; Data curation: VZ, YC; Formal analysis: VZ, YC; Funding acquisition: CU; Investigation: VZ, YC; Methodology: VZ, YC; Project administration: YC, CU; Supervision: YC, CU; Visualization: VZ, YC; Writing-original draft: VZ, YC; Writing-review & editing: VZ, YC, CU.

## Acknowledgement

We thank the JIC Field Experimentation and Horticultural Services teams for their support in glasshouse and field experiments. We thank Dr. Clare Lister on her guidance on lodging testing in the field. We thank Professor Nobuyoshi Watanabe for supplying the germplasm.

## Supplementary Figures

**Supplementary figure 1.**
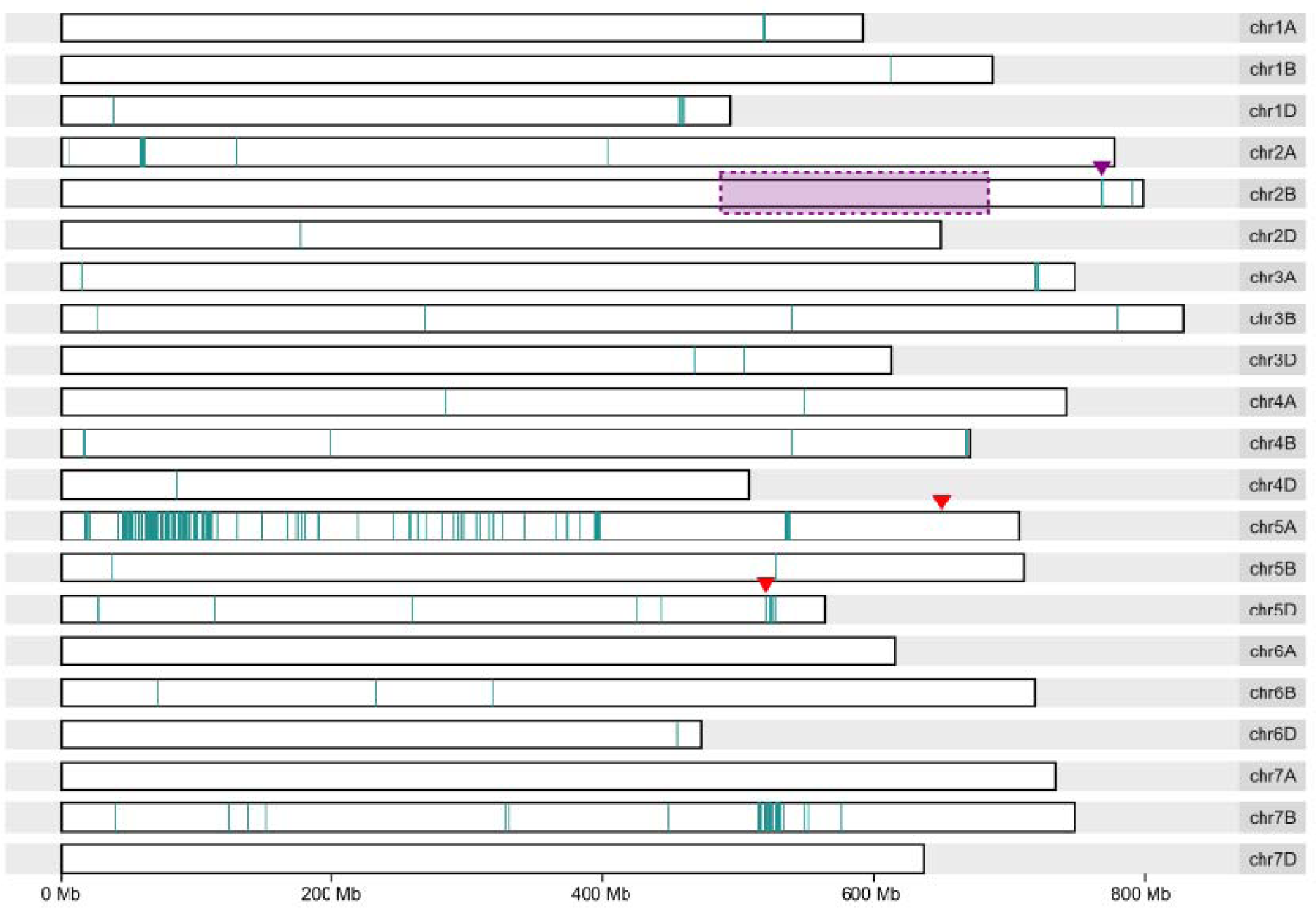
Distribution of SNPs across the 21 chromosomes between N67 and ANK-15 based on RNA-sequencing read alignment to Refseqv1.0. SNPs are represented by vertical bars. The previously mapped location of *C_g_* (Amagai et al., 2016) is indicated by the shaded rectangle, and the polymorphic region closest to this proposed location is indicated by a purple arrowhead. Position of *Q* (*AP2L-A5*; *TraesCS5A02G473800*) and *AP2L-D5* (*TraesCS5D02G486600*) are indicated by red arrowheads.

**Supplementary figure 2.**
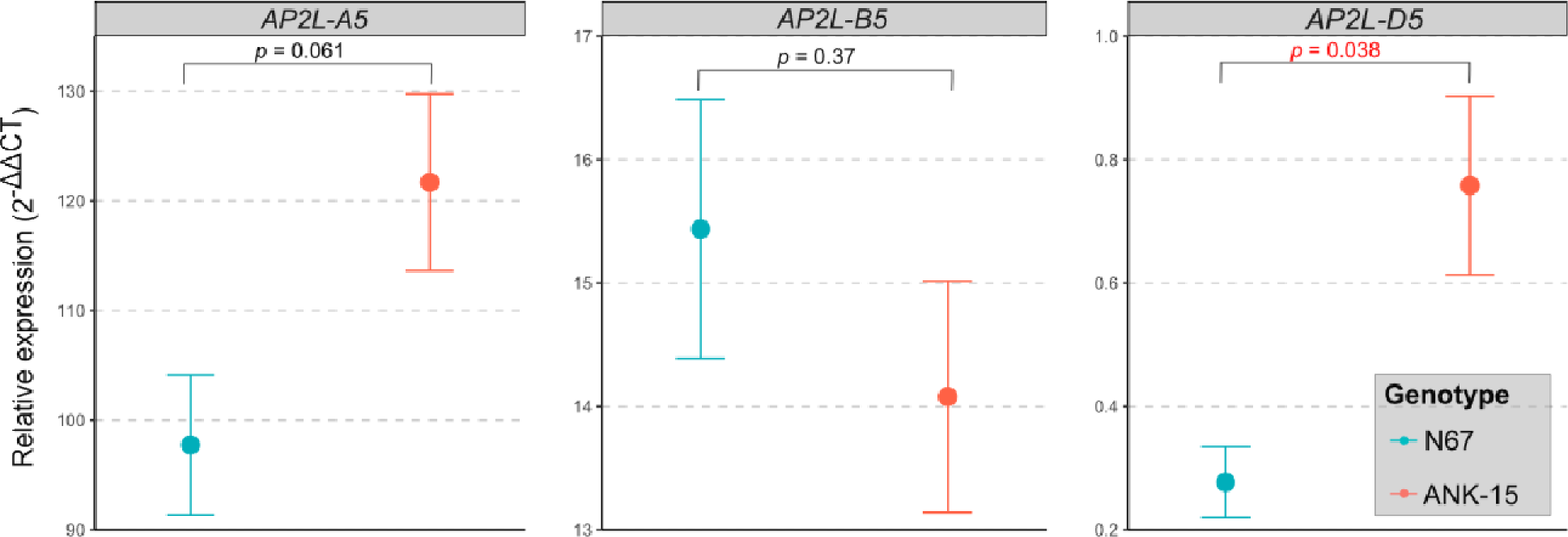
qRT-PCR of *AP2L5* homoeologs in N67 and ANK-15 using rachis collected at yellow anther stage. Points correspond to the mean relative expression of 4 biological replicates, each comprising three technical replicates. Error bars are standard errors of the means. *P-*value is based on Student’s t-test.

**Supplementary figure 3.**
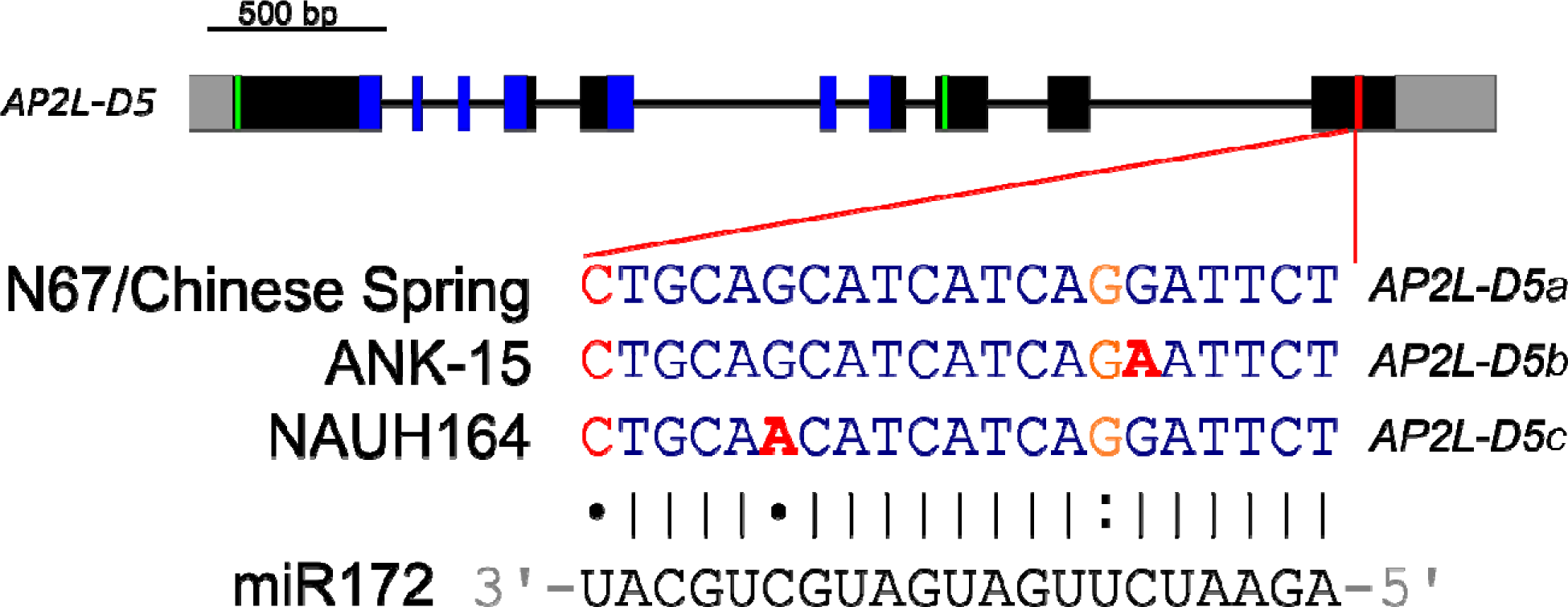
Intron-exon structure of *AP2L-D5* alleles identified in different compact spike head mutants. Both 5’ and 3’ untranslated regions are colored in grey, exons are depicted as black boxes, introns as black lines, AP2 protein domains are highlighted in blue, EAR binding motifs are highlighted in green and the miR172 binding site is highlighted in red. Nucleotides colored red and orange indicate mismatches to miR172 and wobble G:U pairs, respectively. The putative causal SNP leading to the compact spike phenotype of ANK-15 and NAUH164 are shown in bold red font.

**Supplementary figure 4.**
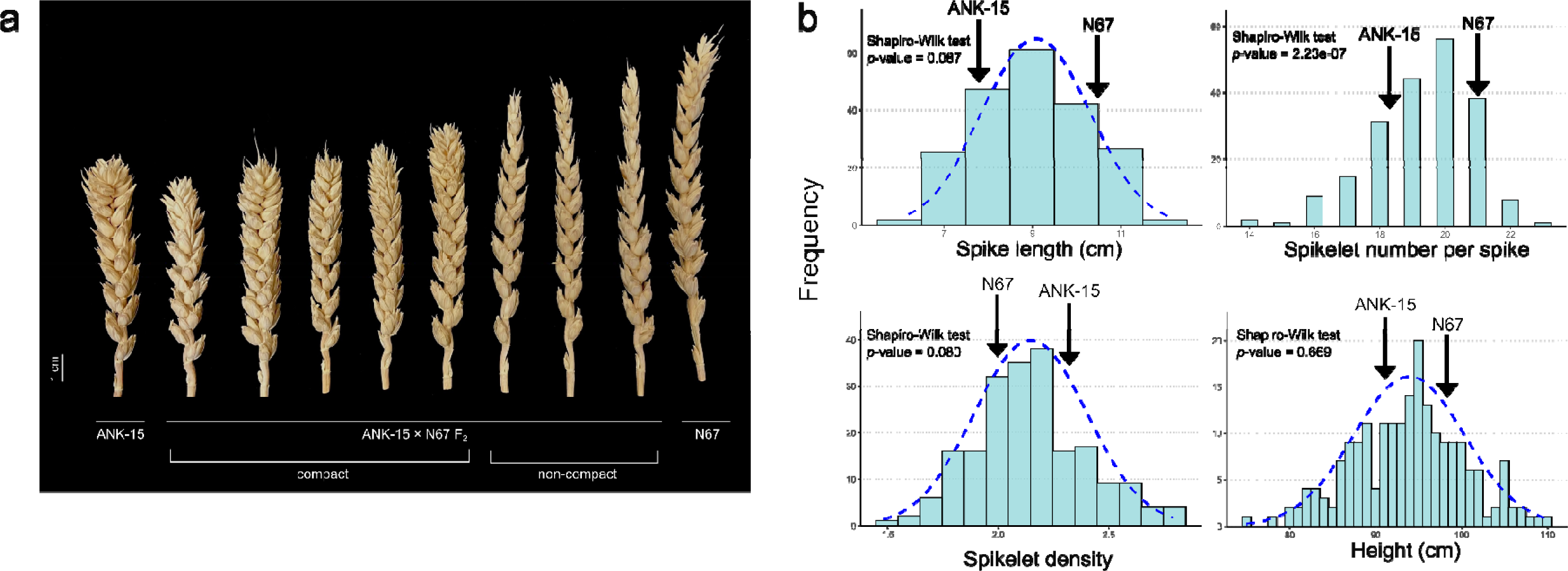
Phenotypic distribution of spike morphology in the N67 × ANK-15 F_2_ mapping population. **a** Variation in spike morphology between N67 and ANK-15 and among the F_2_ population. **b** Frequency distributions of spike length, spikelet number per spike, spikelet density, and height in the F_2_ population. Parental means are indicated with the arrows. Dashed lines represent the normal distribution curve.

**Supplementary figure 5.**
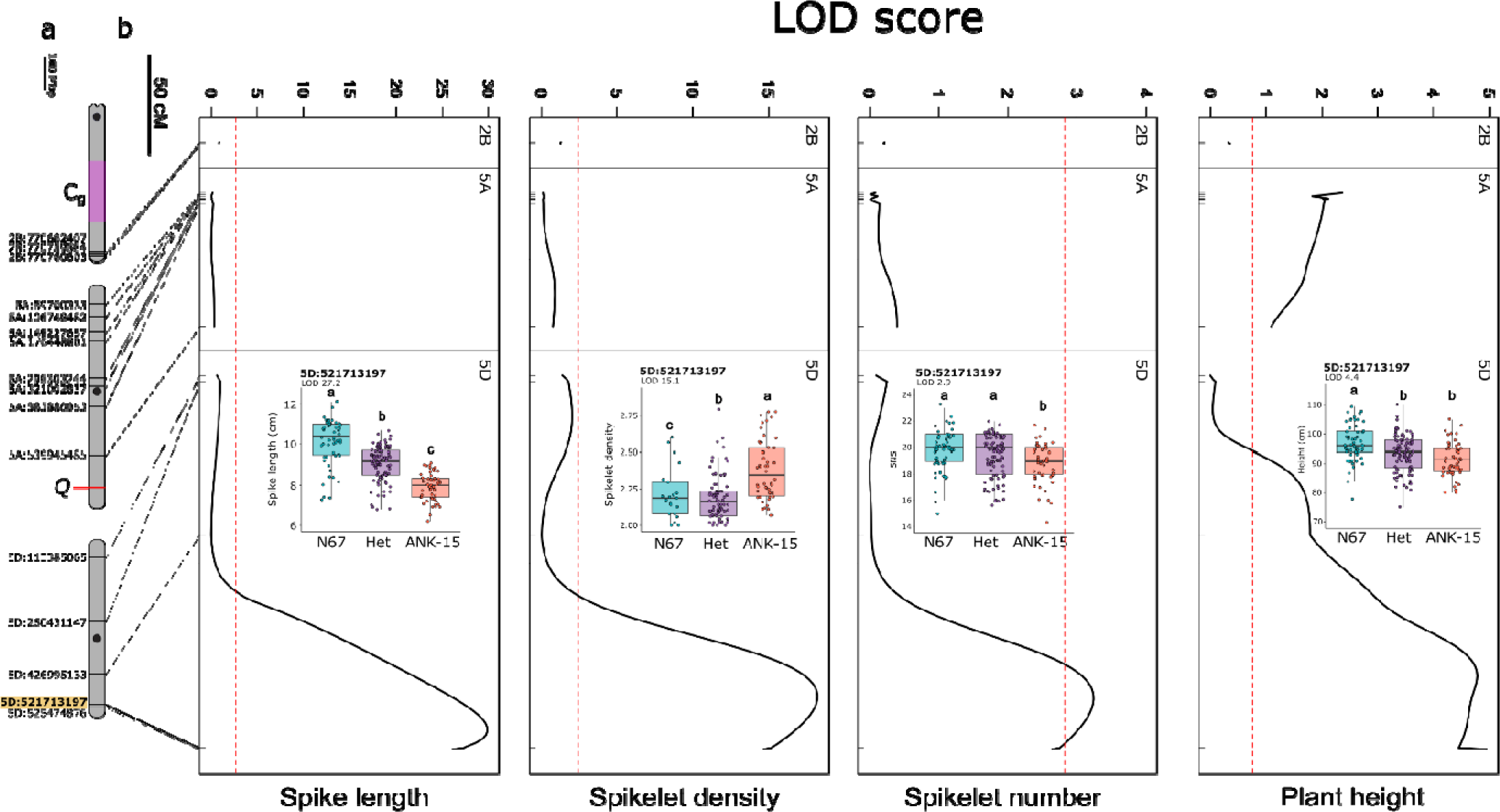
QTL mapping for spike morphology and plant height using N67 × ANK-15 F2 population (n = 207 plants). **a** Physical map of chromosomes 2B, 5A and 5D and the KASP markers designed for genetic mapping. The markers are named after the physical position of the polymorphisms between the parents anchored onto Refseqv1.0 (IWGSC et al. 2018). Black circles denote approximate centromere position. Putative position of the compact spike locus *Cg* is colored in purple. The KASP marker that separates *AP2L-D5a* and *AP2L-D5b* is highlighted in yellow. **b** QTL analysis for spike length, spikelet density, spikelet number and plant height. Genetic positions of the markers are denoted with ticks and connected to their physical locations in a). The dashed red lines denotes the significance threshold estimated based on permutations (n = 1000). **Inset** Spike length distribution of the F_2_ individuals (*n* = 207 plants) carrying parental or heterozygous alleles for KASP marker that separates *AP2L-D5a* and *AP2L-D5b*. Genotype groups significantly different from each other based on post-hoc Tukey test (*P* < 0.01) is denoted with different letters.

## Supplementary Table Heading

**Supplementary table 1. Summary of the read number and the percentage of reads aligned per sample of the RNA-sequencing data**

**Supplementary table 2. High-confidence genes differentially expressed in the rachis tissues between N67 and ANK-15 collected at the green anther stage.**

**Supplementary table 3. Gene ontology terms of the differentially expressed genes in the rachis tissues between N67 and ANK-15 at green anther stage.**

**Supplementary table 4. KASP markers used for QTL analysis of the N67 × ANK-15 F_2_ mapping population (n = 207 plants).**

**Supplementary table 5. List of primers used for sequencing of *AP2L-D5* and qRT-PCR of the *AP2L* homoeologs in wheat.**

