## Supplementary Figure for "Identification of a novel SNP in the miR172 binding site of *Q* homoeolog *AP2L-D5* is associated with spike compactness and agronomic traits in wheat (*Triticum aestivum* L.)"

VZ: 0009-0001-6628-2990

CU: 0000-0002-9814-1770

YC: 0000-0002-8210-1301

^†^ Both authors contributed equally to the work

### Supplementary Figures

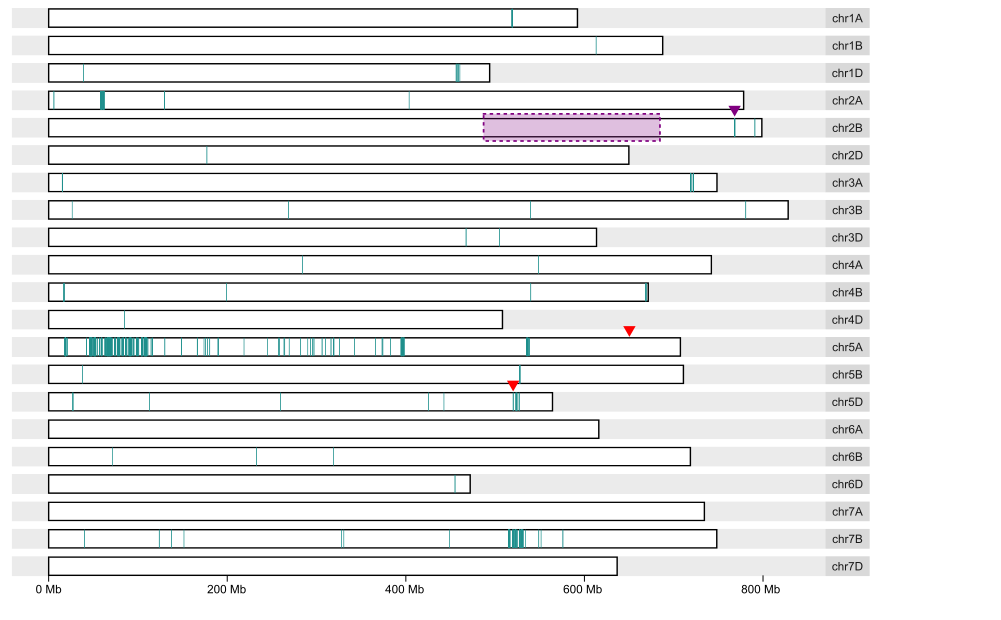

Supplementary figure 1. Distribution of SNPs across the 21 chromosomes between N67 and ANK-15 based on RNA-sequencing read alignment to Refseqv1.0.

SNPs are represented by vertical bars. The previously mapped location of *C_g_* (Amagai et al., 2016) is indicated by the shaded rectangle, and the polymorphic region closest to this proposed location is indicated by a purple arrowhead. Position of *Q* (*AP2L-A5*; *TraesCS5A02G473800*) and *AP2L-D5* (*TraesCS5D02G486600*) are indicated by red arrowheads.

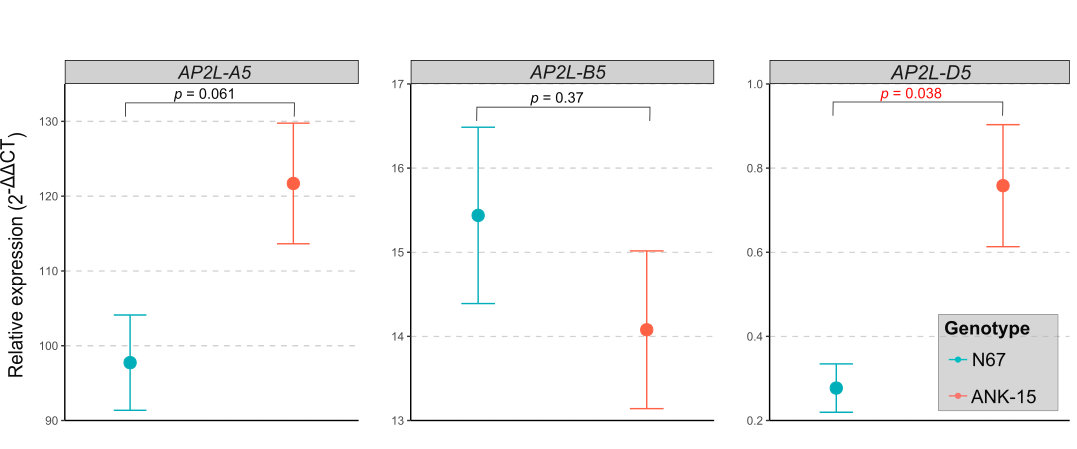

Supplementary figure 2. qRT-PCR of *AP2L5* homoeologs in N67 and ANK-15 using rachis collected at yellow anther stage.

Points correspond to the mean relative expression of 4 biological replicates, each comprising three technical replicates. Error bars are standard errors of the means. *P-*value is based on Student’s t-test.

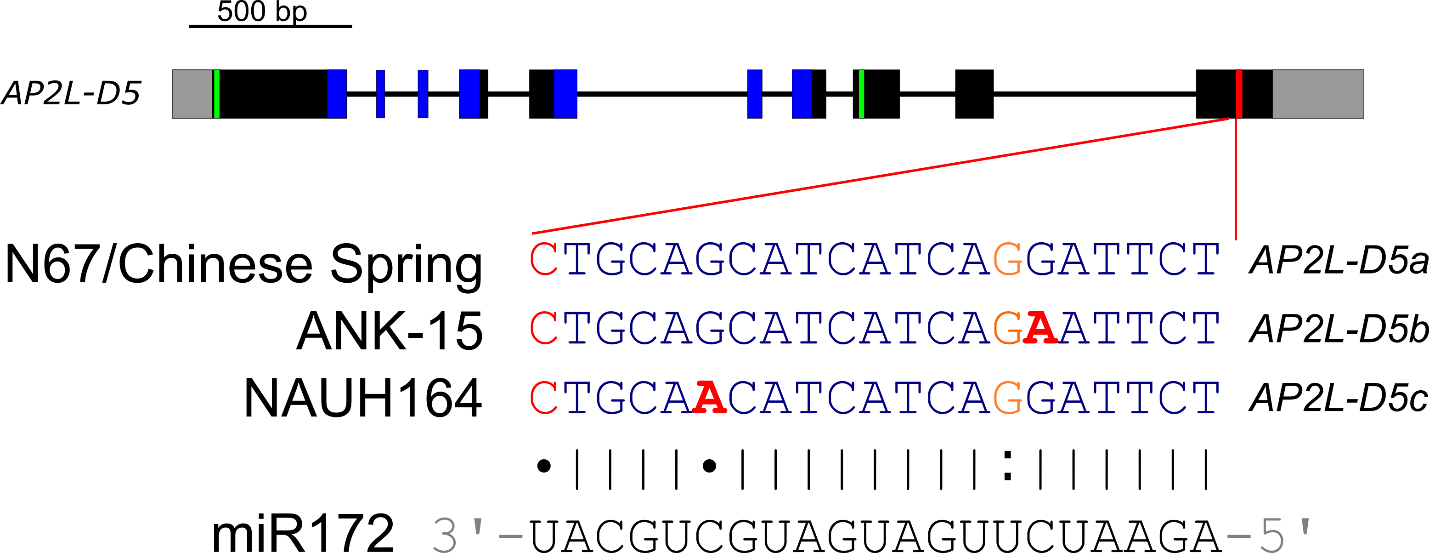

Supplementary figure 3. Intron-exon structure of *AP2L-D5* alleles identified in different compact spike head mutants.

Both 5’ and 3’ untranslated regions are colored in grey, exons are depicted as black boxes, introns as black lines, AP2 protein domains are highlighted in blue, EAR binding motifs are highlighted in green and the miR172 binding site is highlighted in red. Nucleotides colored red and orange indicate mismatches to miR172 and wobble G:U pairs, respectively. The putative causal SNP leading to the compact spike phenotype of ANK-15 and NAUH164 are shown in bold red font.

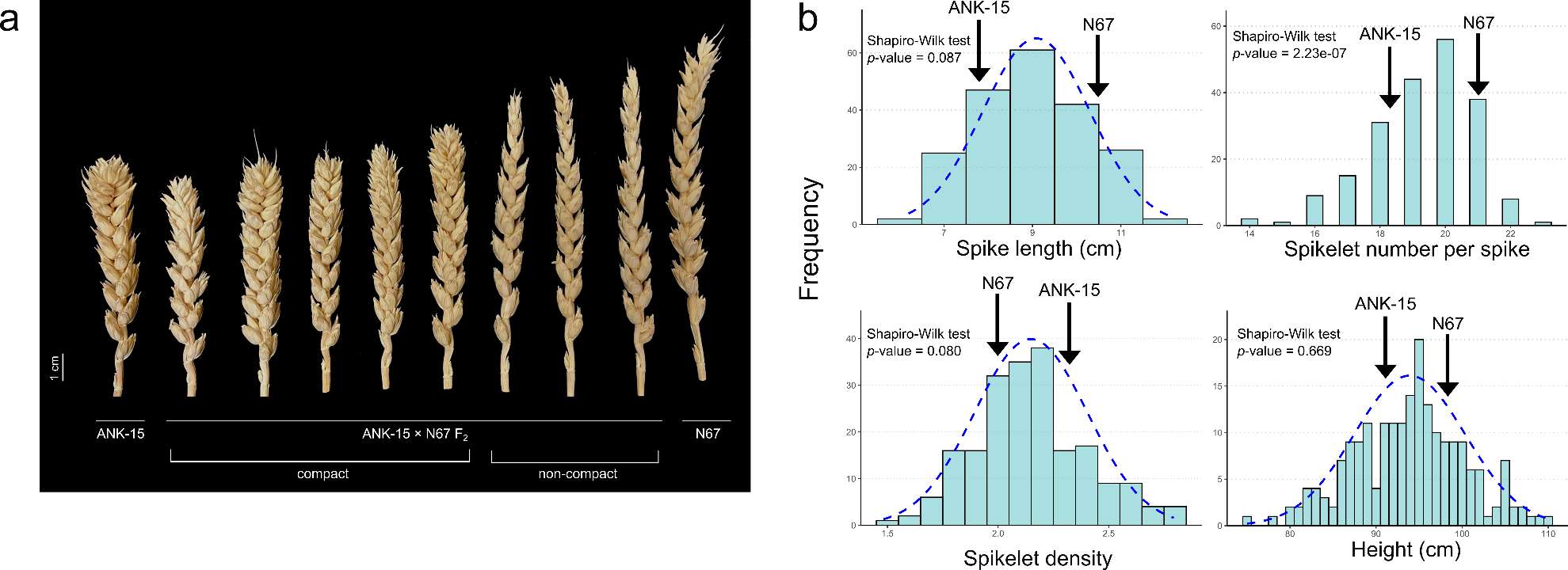

Supplementary figure 4. Phenotypic distribution of spike morphology in the N67 × ANK-15 F_2_ mapping population.

**a** Variation in spike morphology between N67 and ANK-15 and among the F_2_ population. **b** Frequency distributions of spike length, spikelet number per spike, spikelet density, and height in the F_2_ population. Parental means are indicated with the arrows. Dashed lines represent the normal distribution curve.

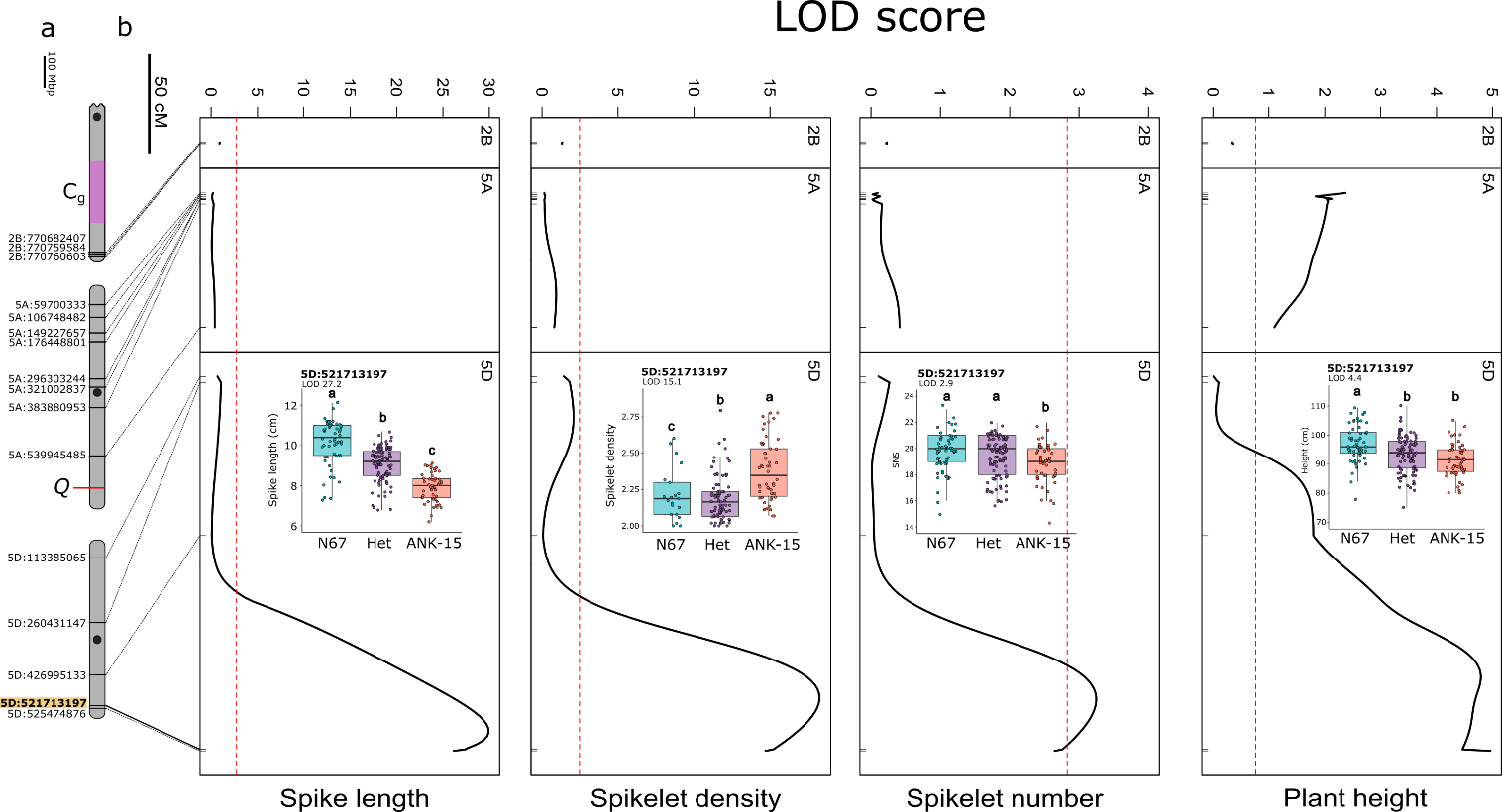

Supplementary figure 5. QTL mapping for spike morphology and plant height using N67 × ANK-15 F_2_ population (n = 207 plants).

**a** Physical map of chromosomes 2B, 5A and 5D and the KASP markers designed for genetic mapping. The markers are named after the physical position of the polymorphisms between the parents anchored onto Refseqv1.0 (IWGSC et al. 2018). Black circles denote approximate centromere position. Putative position of the compact spike locus *Cg* is colored in purple. The KASP marker that separates *AP2L-D5a* and *AP2L-D5b* is highlighted in yellow. **b** QTL analysis for spike length, spikelet density, spikelet number and plant height. Genetic positions of the markers are denoted with ticks and connected to their physical locations in a). The dashed red lines denotes the significance threshold estimated based on permutations (n = 1000). **Inset** Spike length distribution of the F_2_ individuals (*n* = 207 plants) carrying parental or heterozygous alleles for KASP marker that separates *AP2L-D5a* and *AP2L-D5b*. Genotype groups significantly different from each other based on post-hoc Tukey test (*P* < 0.01) is denoted with different letters.
